# Metagenomes and Metagenome-Assembled Genomes from Microbial Communities in a Biological Nutrient Removal Plant Operated at Los Angeles County Sanitation District (LACSD) with High and Low Dissolved Oxygen Conditions

**DOI:** 10.1101/2025.11.10.687646

**Authors:** Blaise M. Enuh, Kevin S. Myers, Phil Ackman, Thomas Weiland, Natalie Beach, Michelle Young, Timothy J. Donohue, Daniel R. Noguera

## Abstract

Aeration represents one of the largest energy costs in water resource recovery facilities (WRRFs). Previous studies have shown that effective nitrification and phosphorus removal can be maintained in biological nutrient removal (BNR) systems operated under low dissolved oxygen (DO) conditions, significantly reducing energy use. To improve understanding of microbial community adaptation to reduced DO, we analyzed metagenomes and metagenome-assembled genomes (MAGs) from the Pomona WRRF (Los Angeles County Sanitation Districts) before and after a gradual decrease in operating DO from approximately 3.5 mg/L to 0.7 mg/L over 18 months. Metagenomic DNA was isolated from high and low DO samples, then sequenced using PacBio HiFi technology. A total of 492 MAGs were recovered of which 304 were unique after dereplication. These genomes expand current knowledge of microbial community dynamics in low-DO BNR systems and provide valuable genomic resources for understanding microbial adaptation to energy-efficient wastewater treatment processes.

## Background

Aeration at water resource recovery facilities (WRRF) can be a major operational expense. It has been demonstrated that to reduce electricity use, effective nitrification and phosphorus removal can be achieved in biological nutrient removal (BNR) operated with low dissolved oxygen (DO) levels (Fitzgerald et al., 2015; Keene et al., 2017; Stewart et al., 2021). The microbial community adapts to the altered environmental conditions when DO is reduced in BNR plants (Fitzgerald et al., 2015; Park and Noguera, 2008, 2004). To contribute to the understanding of microbial community changes that occur upon lowering DO in a full-scale process, we report here metagenomes and metagenome-assembled genomes (MAGs) from the Pomona WRRF operated by the Los Angeles County Sanitation Districts (LACSD) before and after an experiment that reduced operating DO from approximately 3.5 mg/L to 0.70 mg/L in 18 months (Carollo Engineers et al., 2025). Five samples were analyzed, three from high-DO operation and two from low-DO operation.

## Metagenomic DNA extraction, library preparation, sequencing, and analysis

DNA isolation was performed using the DNeasy PowerSoil Kit using the published protocol (Qiagen, Germantown, MD) and quantified via Qubit fluorometer (Fisher Scientific, Waltham, MA), then stored at −20°C prior to sequencing. DNA purity was assessed using a NanoDrop One spectrophotometer (Fisher Scientific). DNA concentration was determined with the Qubit dsDNA High Sensitivity assay following standard protocols (Fisher Scientific).

Samples were processed at the University of Wisconsin-Madison Biotechnology Center (Madison, WI) for library construction and sequencing. HiFi libraries were generated following protocol PN 102-166-600 Version 04 (Pacific Biosciences, Menlo Park, CA) with standard parameters. The procedure involved fragmentation with g-TUBEs (Covaris, Woburn, MA) according to the shearing protocol for large genomes (2,164 x g) and size selection using diluted AMPurePB beads (Pacific Biosciences) as detailed in the procedure-checklist. Library integrity was evaluated on the FemtoPulse System (Agilent, Santa Clara, CA). Quantification was performed using the Qubit dsDNA High Sensitivity assay, followed by sequencing on a Sequel II platform with Sequel Polymerase Binding Kit 2.2 following the standard protocol (Pacific Biosciences). Default parameters for all programs were used unless otherwise stated. Sequencing was done separately at High and low DO sampling time points. The resulting Circular Consensus Sequence (CCS) reads were then assembled utilizing metaFLYE (v2.9-b1768) (Kolmogorov et al., 2020) and metaMDBG (v0.3) (Benoit et al., 2024) then polished with racon (v1.4.20) (Vaser et al., 2017). Subsequently, the reads were mapped onto the assemblies using minimap2 (v2.22-r1101) (Li, 2018). Assembly binning was done with metaBAT2 (v2:2.15) (Kang et al., 2019). Contaminated contigs were identified using ProDeGe (v2.3) (Tennessen et al., 2016) and custom scripts for tetranucleotide frequency analysis (run.GC.sh and Calculating_TF_Correlations.R; https://github.com/GLBRC/metagenome_analysis). Contigs flagged by both methods were removed from the final MAG assemblies. All MAGs were evaluated for quality using CheckM (v1.2.2) (Parks et al., 2015) and the taxonomy of each MAG was determined using GTDB-Tk (v2.1.0) database release 09-RS214 (Chaumeil et al., 2020). Functional annotations for each MAG were assigned using Bakta (v1.9.1) (Schwengers et al., 2021).

RAxML-NG adaptive (v 1.2.1) (Kozlov et al., 2019) with default parameters and the multiple sequence alignment of 120 bacterial marker genes generated by GTDB-Tk was used for maximum likelihood-based inference of the best phylogenetic tree. The resulting phylogenetic tree was visualized and further annotated using TreeViewer (v2.2.0)(Bianchini and Sánchez-Baracaldo, 2024) and Inkscape (v1.2.2) (The Inkscape team, 2025). The final tree is shown in Figure 1.

**Figure 1.**
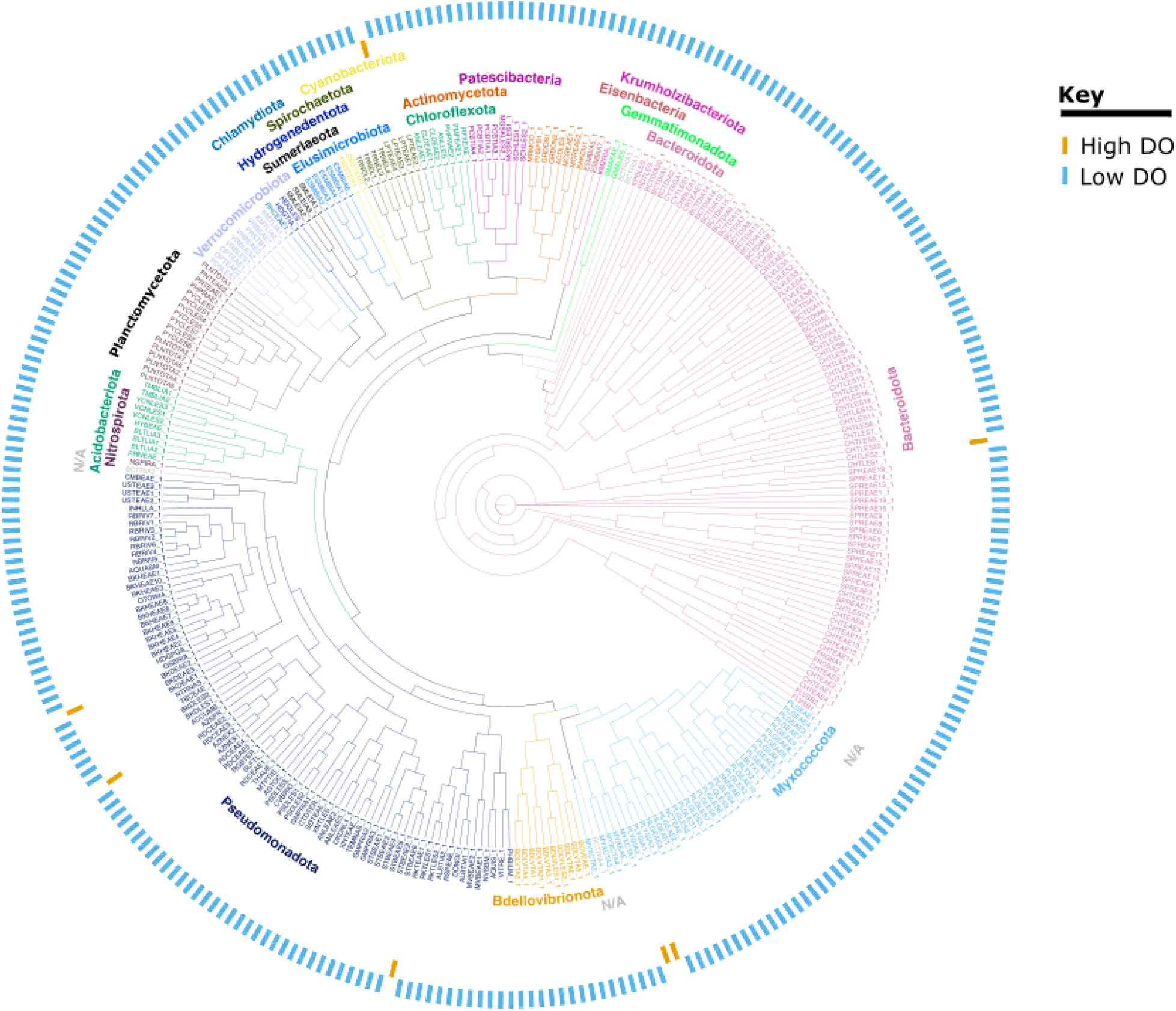
Phylogenetic map showing an overview of community structure based on the 304 unique MAGs from the LACSD plant assembled from both low and high DO samples. The suffix ‘_1’ was assigned to MAGs determined by dRep to be the best in their respective clusters. These are the representatives that were used to generate the plot. From the innermost to the outermost ring: The innermost ring represents phylogenetic clustering of MAGs. The names of each phylum are colored according to the clusters they represent. N/A indicates MAGs that were not classified into any phylum. The next ring in orange points to MAGs that were assembled from the high DO samples while the blue ring immediately after it points to MAGs that were assembled from the low DO samples.

In total, we obtained 463 MAGs from the low DO samples and 29 from the high DO samples (Supplementary File S1). Of these, 304 were deemed to be unique after dereplication with dRep (v0.6.1) (Olm et al., 2017) (Figure 1). These data augment the knowledge base of microbial communities in BNR operated at low DO. All the MAGs and corresponding details about their quality, classification, and genome characteristics are provided in the Supplementary File S1.

## Acknowledgments

This work was supported by funding from the U.S. Department of Energy (DOE), Office of Energy Efficiency & Renewable Energy, under award DE-EE0009509, and partially based upon work at the Great Lakes Bioenergy Research Center supported by the U.S. Department of Energy, Office of Science, Biological and Environmental Research under Award Number DE-SC0018409. DNA sequencing was carried out at the UW Biotechnology Center’s DNA Sequencing Facility (Research Resource Identifier – RRID:SCR_017759).

## Data availability

Fastq files for the metagenomes have been deposited in the NCBI SRA database under Bio project number PRJNA1333415. The MAGs and their annotation dataset can be accessed on Figshare (10.6084/m9.figshare.30412474). The custom scripts used in these analyses are available at the Great Lakes Bioenergy Research Center GitHub repository (https://github.com/GLBRC/metagenome_analysis).

